# A bispecific antibody-drug conjugate targeting pCAD and CDH17 has antitumor activity and improved tumor specificity

**DOI:** 10.1101/2024.04.19.590291

**Authors:** Alyssa Synan, Nila C Wu, Roberto Velazquez, Claude Logel, Kathrin Mueller, Andrew Green, Patrizia Barzaghi-Rinaudo, Quincey Simmons, Samuele Mercan, Xingyi Shi, Joshua Korn, Margaret McLaughlin, William R Tschantz, Dominik Hainzl, Anthony Malamas, Regis Cebe, Kathleen T Xie, Joseph A D’Alessio

**Author notes:** Co-corresponding authors: Kathleen T Xie, Joseph A D’Alessio. These authors contributed equally (Alyssa Synan, Nila C Wu).

## Abstract

P-cadherin (pCAD) and LI-cadherin (CDH17) are cell surface proteins belonging to the cadherin superfamily that are both highly expressed in colorectal cancer. This co-expression profile presents a novel and attractive opportunity for a dual targeting approach using an antibody-drug conjugate (ADC). In this study, we used a unique avidity-driven *in vitro* screening approach to generate pCAD x CDH17 bispecific antibodies that selectively targets cells expressing both antigens over cells expressing only pCAD or only CDH17. Based off the *in vitro* results we selected a lead bispecific antibody to link to the cytotoxic payload MMAE to generate a pCAD x CDH17 bispecific MMAE ADC. In *in vivo* dual flank mouse models, we demonstrated antitumor activity of the bispecific ADC in tumors expressing both antigens, but not in tumors expressing only pCAD or only CDH17. Overall, the preclinical data presented here suggests that a pCAD x CDH17 bispecific MMAE ADC has the potential to provide clinical benefit to colorectal cancer patients.

## Introduction

Antibody-drug conjugates (ADCs) are an innovative class of targeted therapeutics that have the potential for improved cancer treatment. ADCs combine the selectivity of monoclonal antibodies with the potent cytotoxic activity of small molecule drugs (payloads), enabling targeted payload delivery through antigen-specific binding. To induce cell cytotoxicity, ADCs bind to the surface antigen target, internalize within the cell, and subsequently traffic to the lysosome where the antibody-payload linker or antibody is degraded, releasing the cytotoxic payload to act on its intended target^1,2^. Despite the promise of ADCs, there remain challenges in their development and clinical implementation, such as achieving sufficient tumor specificity that minimizes damage to normal tissues induced by the highly potent payload^3,4^.

P-cadherin (pCAD, placental Cadherin, Cadherin-3; encoded by CDH3) is a cell surface glycoprotein and a member of the classical cadherin superfamily. It is involved in calcium-dependent cell-cell adhesions of the epithelium and is associated with various tumor-promoting processes, including cell invasiveness, metastasis, and tumorigenesis^5,6^. pCAD is overexpressed in several cancer types while being restricted in normal tissues^6^, presenting an attractive antigen for delivering potent cytotoxic payloads. Indeed, PCA062, a first-in-class pCAD-targeting ADC, demonstrated potent *in vitro* and *in vivo* antitumor activity in several pCAD-expressing tumors and achieved a favorable safety profile in non-human primate toxicology studies^7^.

CDH17 (LI-Cadherin, liver-intestinal Cadherin, Cadherin-17; encoded by CDH17) is similarly part of the cadherin superfamily and plays an essential role in cell adhesion and organ development, particularly in the formation and maintenance of the intestinal epithelium^8,9^. In colorectal cancer (CRC), CDH17 has been implicated in regulating integrin signaling for cell adhesion, promoting cell proliferation, inhibiting apoptosis and facilitating metastasis^10,11^. Its expression has been associated with prognostic importance and has therefore emerged as a key CRC marker^12^. While various CDH17-based therapeutics are under investigation for treating CRC including an antibody^13^, an antibody-conjugated to a photosensitizer^14^, and a T-cell engager^15^, there are no CDH17-targeted ADCs that have reached clinical trials. Notably, pCAD and CDH17 are both highly expressed in CRC, presenting an opportunity to leverage their combination to improve ADC specificity to CRC cells expressing both antigens.

Bispecific antibodies (BsAbs) are designed to simultaneously target two distinct antigens. Previous studies combining BsAbs with the ADC approach have demonstrated advantages, including enhanced ADC efficacy through improved internalization and lysosome-directed degradation. For example, coupling the rapidly internalizing protein PRLR with the clinically approved ADC target HER2 was shown to enhance internalization and degradation of HER2, resulting in increased cell death compared to a monospecific HER2 ADC^16^ (Andreev, 2017). Similarly, coupling the lysosomal membrane protein CD63 with HER2 induced ADC lysosomal accumulation in HER2+ cells, leading to increased cell cytotoxicity^17^ (de Goeij, 2017). Other bispecific ADCs have been developed that couple the tyrosine kinase receptor EGFR with targets such as cMET, aiming to overcome secondary pathway mutations that cause treatment resistance^18^, or with MUC1^19^ or TROP2^20^, which are highly co-expressed on cancer tissues. Notably, the EGFR x cMET ADC and the EGFR x MUC1 ADC are currently in clinical trials.

The objective of our work was to enhance ADC tumor specific potency through the required engagement of multiple antigens using a pCAD x CDH17 BsAb, broadening the therapeutic index by limiting on-target, off-tumor toxicity on healthy normal cells expressing either pCAD or CDH17 only. Towards identifying a lead pCAD x CDH17 bispecific ADC, we designed a unique and novel screening approach where we screened CDH17 arms and affinity-tuned pCAD arms based on avidity. We utilized *in vitro* cancer cell lines and assays that assess BsAb binding, and internalization to direct the cytotoxic molecule MMAE, to dual-antigen expressing cells compared to single antigen-expressing cells. The whole screening approach was conducted in two phases. First, we screened CDH17 arms and selected a high avidity candidate as the lead, based on its ability to produce the greatest difference between a one-armed (monovalent) antibody and its corresponding full-length (bivalent) antibody for cell binding and inhibiting cell proliferation. Next, we generated a panel of affinity-tuned pCAD arms and similarly screened for *in vitro* cell cytotoxicity and binding differentials. Through this rationally designed two-phase workflow, we identified a lead pCAD x CDH17 BsAb, which we subsequently conjugated with MMAE payload. In preclinical *in vivo* studies using mice bearing dual flank tumors of CRC, we demonstrated that the pCAD x CDH17 bispecific ADC inhibited the growth of dual-antigen expressing tumors while sparing single-antigen expressing tumors. Overall, these results support the development of bispecific ADCs and validate pCAD x CDH17 as paired targets, with the potential to improve the therapeutic index and offer clinical benefit to cancer patients.

## Results

### pCAD and CDH17 dual target expression is high in solid tumors and limited in normal tissues

To validate the suitability of the targets for a bispecific ADC approach, we first set out to evaluate the bulk mRNA levels of *CDH3* (the gene encoding pCAD) and *CDH17* in both human normal tissue and cancer samples using publicly available datasets from The Cancer Genome Atlas (TCGA)^21^ and Gene-Tissue-Expression (GTEX)^22^. Our analysis revealed that *CDH3* mRNA is overexpressed in various solid tumors, including CRC, and found at moderate levels in normal esophagus, skin, prostate, and reproductive tissues **(Figure 1Ai)**. *CDH17* mRNA is similarly overexpressed in several gastrointestinal solid tumors, including pancreas and stomach cancer in addition to CRC. In normal tissues, *CDH17* mRNA is moderately expressed in the large and small intestine **(Figure 1Aii)**. These findings are also reflected in publicly available scRNAseq datasets for healthy normal donors from Tabula Sapiens^23^, where *CDH3* single cell RNA expression is widespread across tissues, and *CDH17* expression is mostly expressed in the small and large intestine **(Figure S1A)**. We also integrated multiple publicly available scRNAseq datasets^24,25^ collected from CRC patients to confirm that cancer cells are the cell type responsible for high *CDH17* and *CDH3* expression, instead of other cell types present in the CRC tumor microenvironment **(Figure S1B)**. While there are detectable mRNA levels of *CDH3* and *CDH17* in specific normal tissues independently, for the applicability of the dual-targeting approach we were most interested in evaluating co-expression of *CDH3* and *CDH17* across normal tissue and CRC samples. Encouragingly, our co-expression analysis demonstrated that the *CDH3* and *CDH17* dual expressing population is predominantly isolated to cancer cells, with minimal dual-positive expression observed in normal tissues **(Figure 1B)**.

**Figure 1.**
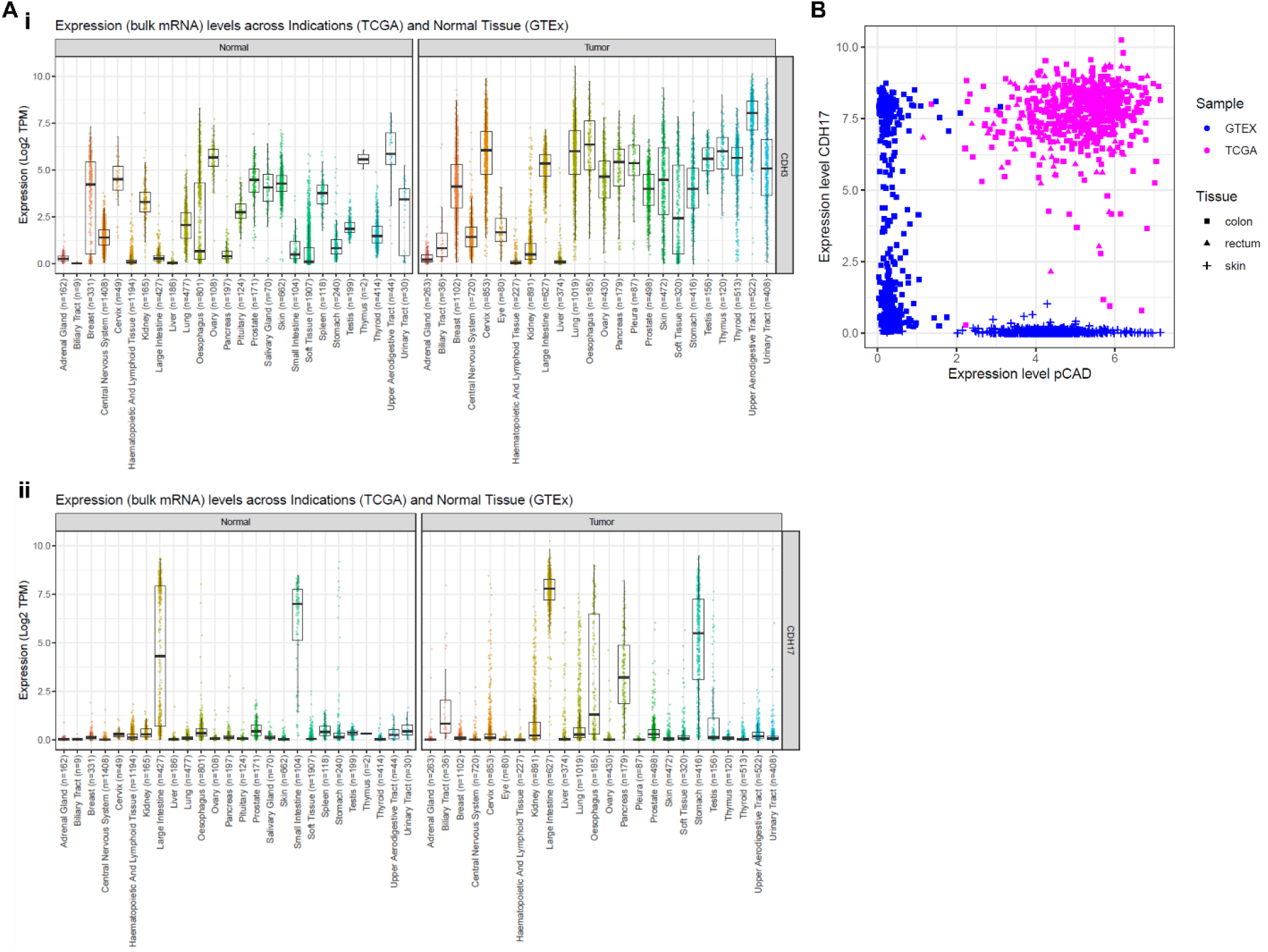
pCAD and CDH17 target expression are high in solid tumors and limited in normal tissues. (A) Bulk RNA expression of (i) *CDH3* and (ii) *CDH17* across cancer types and normal tissue samples obtained from TCGA and GTEx databases, respectively. (B) Correlation between *CDH17* and *CDH3* expression in CRC and normal tissue samples.

### Generation of BsAb phase I: selection of lead CDH17 arm

Based on the suitability of pCAD and CDH17 as antigens for a dual antigen targeting approach to increase tumor-targeting selectivity, we next set out to generate a cross-arm avidity binder following the schematic outlined in **Figure 2A**.

**Figure 2.**
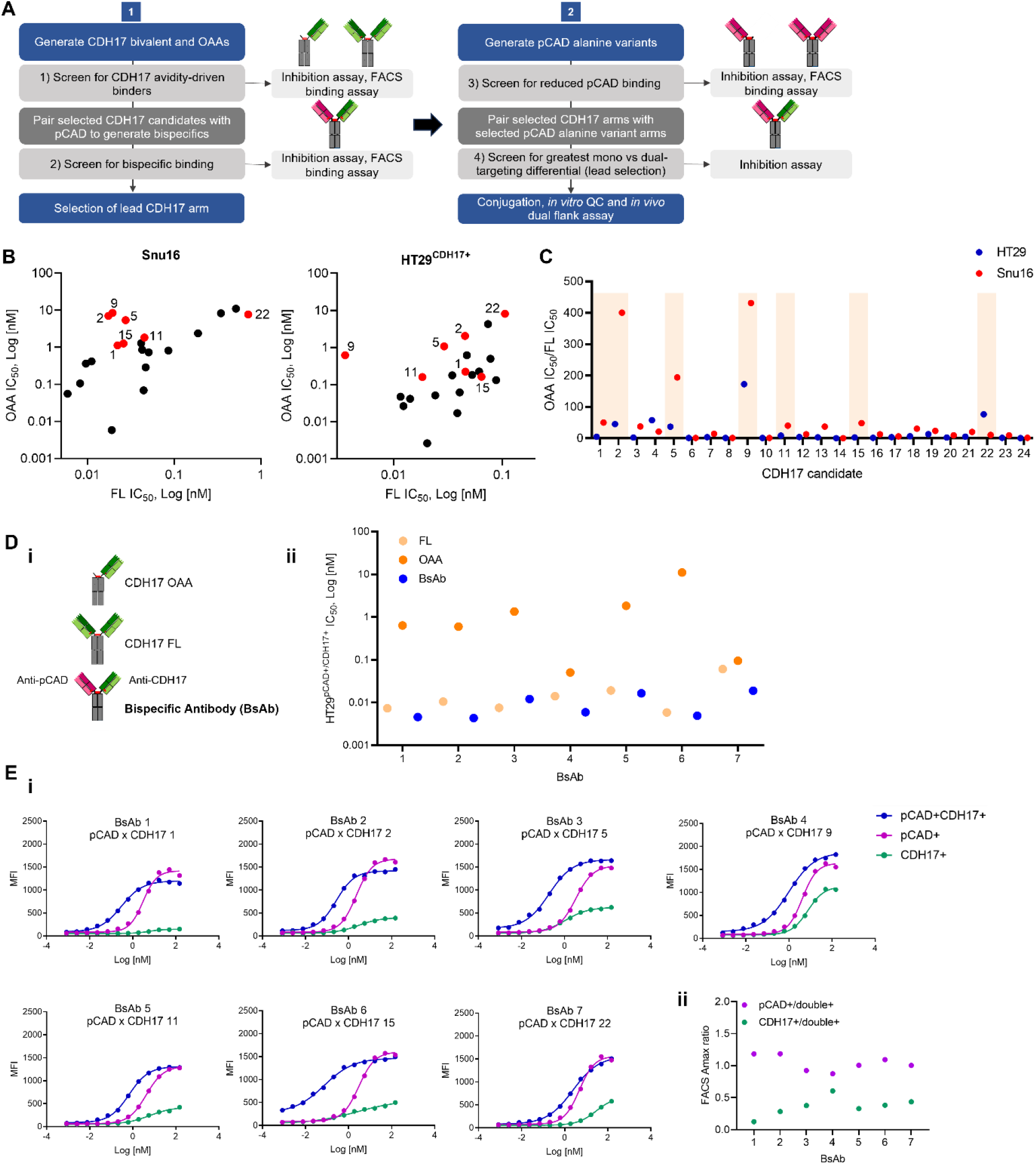
Selection of CDH17 arm and generation of 1st round of bispecifics. (A) A schematic outlining the strategy for the identification of a lead pCAD x CDH17 BsAb. The first phase involves screening CDH17 monovalent and bivalent antibodies, and pairing the most avid CDH17 arm with a pCAD arm to evaluate potency and binding. The second phase involves screening a panel of pCAD alanine variants to identify the most avid pCAD arm, followed by pairing with the lead CDH17 arm. (B) One-armed antibody (OAA, monovalent) vs full-length antibody (FL, bivalent) internalization and growth inhibition results for the panel of CDH17 candidates in Snu16 and HT29^CDH17+^ cell lines. A red dot indicates the top 7 CDH17 candidates. (C) The IC_50_ fold changes between the OAA and FL antibody in both cell lines, with the top candidates highlighted. (D) (i) The top CDH17 candidates were paired with an anti-pCAD arm (generating BsAb 1-7) and compared to the corresponding CDH17 OAA and FL CDH17 arms. (ii) Internalization and proliferation inhibition assay in HT29^pCAD+CDH17+^ treated with the three formats described in (i). (E) (i) FACS binding titration data of the 7 BsAbs in HT29^pCAD+^, HT29^CDH17+^ HT29^pCAD+CDH17+^ demonstrated that the BsAbs preferentially bind to dual expressing cells compared to CDH17+ cells. (ii) Fold change of the Amax of the BsAb binding curves between the pCAD+ and double positive cell or CDH17+ and double positive cell.

The first phase of the selection process involved identifying an avidity driven CDH17 binding arm. We generated 24 unique anti-CDH17 IgG antibodies to be functionally profiled. To enable screening for high avidity binders, we also generated 24 one-armed antibodies (OAA) to be compared to the corresponding 24 full length (FL) CDH17 antibodies. First, we evaluated the 48 total OAA and FL candidates using an *in vitro* internalization and inhibition of cell proliferation assay in two CDH17+ cell lines, Snu16 (stomach cancer cell line), and an engineered CDH17-overexpressing CRC cell line, HT29^CDH17+^ **(Figure 2B, Figure S2A)**. Our analysis demonstrated that IC_50_ values ranged from 1.8E-6 nM to 10.9 nM in Snu16 cells, and 8.4E-6 nM to 11 nM in HT29^CDH17+^ cells. To enable selection based on highly avid arms, we calculated the ratio between IC50 values of corresponding OAA and FL antibodies and filtered the list of candidates based on those with a high ratio observed across both cell lines tested **(Figure 2C)**. The resulting top 7 CDH17 arms were: CDH17 1, CDH17 2, CDH17 5, CDH17 9, CDH17 11, CDH17 15, and CDH17 22, as highlighted in **Figure 2C**. The 48 total antibodies were also assessed in a FACS binding assay in HT29^CDH17+^ cells, and the ratio between binding IC50s of corresponding OAA and FL antibodies were similarly quantified **(Figure S3A)**. Across the 7 CDH17 arms selected from the *in vitro* internalization and inhibition of proliferation assay, all OAA antibodies had lower or similar binding to HT29^CDH17+^ cells compared to the respective FL antibodies **(Figure S3B)**.

Upon filtering the 24 candidates, we next combined all 7 CDH17 arms with a pCAD arm to generate 7 pCAD x CDH17 BsAbs, named BsAb 1 through 7 **(Figure 2Di)**. To evaluate whether the avid property of the CDH17 arm was maintained in the bispecific format, we compared the internalization and proliferation inhibition capacity between corresponding OAAs, FL antibodies, and BsAbs, in an engineered dual-expressing CRC cell line, HT29^pCAD+CDH17+^ **(Figure 2Dii)**. Our results demonstrated that all 7 BsAbs retained higher efficacy at internalizing and inhibiting proliferation compared to their corresponding OAAs, with similar potency to their FL equivalents. The potency of our BsAbs was further translated to dual-positive cell selectivity, upon assessing the cell binding of the 7 BsAbs in single positive expressing cell lines (HT29^pCAD+^ and HT29^CDH17+^) compared to a double positive expressing cell line (HT29^pCAD+CDH17+^) **(Figure 2Ei)**. All 7 BsAbs showed increased dual antigen specific binding compared to the CDH17-only expressing cell line, quantified by the fold change between their measured Amax values **(Figure 2Eii)**. While we achieved a lack of selectivity for CDH17-only expressing cells using our avidity screening approach, the difference between the dual expressing cell line and pCAD-only expressing cell line was minimal, justifying the search for an avidity-driven pCAD arm. Before the subsequent phase of our approach however, we further narrowed down our 7 CDH17 candidates to a lead CDH17 arm that had the most avid arm, CDH17 11, based off the combined internalization and proliferation inhibition and FACS binding assay results. Overall, in the first pCAD x CDH17 bsAb search phase, we demonstrated improved selectivity of our lead CDH17 arm to dual expressing cells compared to CDH17-only expressing cells.

### Generation of BsAb phase II: selection of lead affinity detuned pCAD arm

The second phase of the selection process set out to address the remaining activity on pCAD-only expressing cells, in an attempt to mitigate any potential toxicity to pCAD normal tissue liabilities as previously discussed. To generate additional pCAD candidates for testing, we detuned the affinity of the pCAD arm by alanine scanning the light chain CDR3, where each amino acid of the CDR was systematically replaced by an alanine, generating a panel of 8 light chain single alanine variants named pCAD var 1 through 8 **(Figure 3A)**.

**Figure 3.**
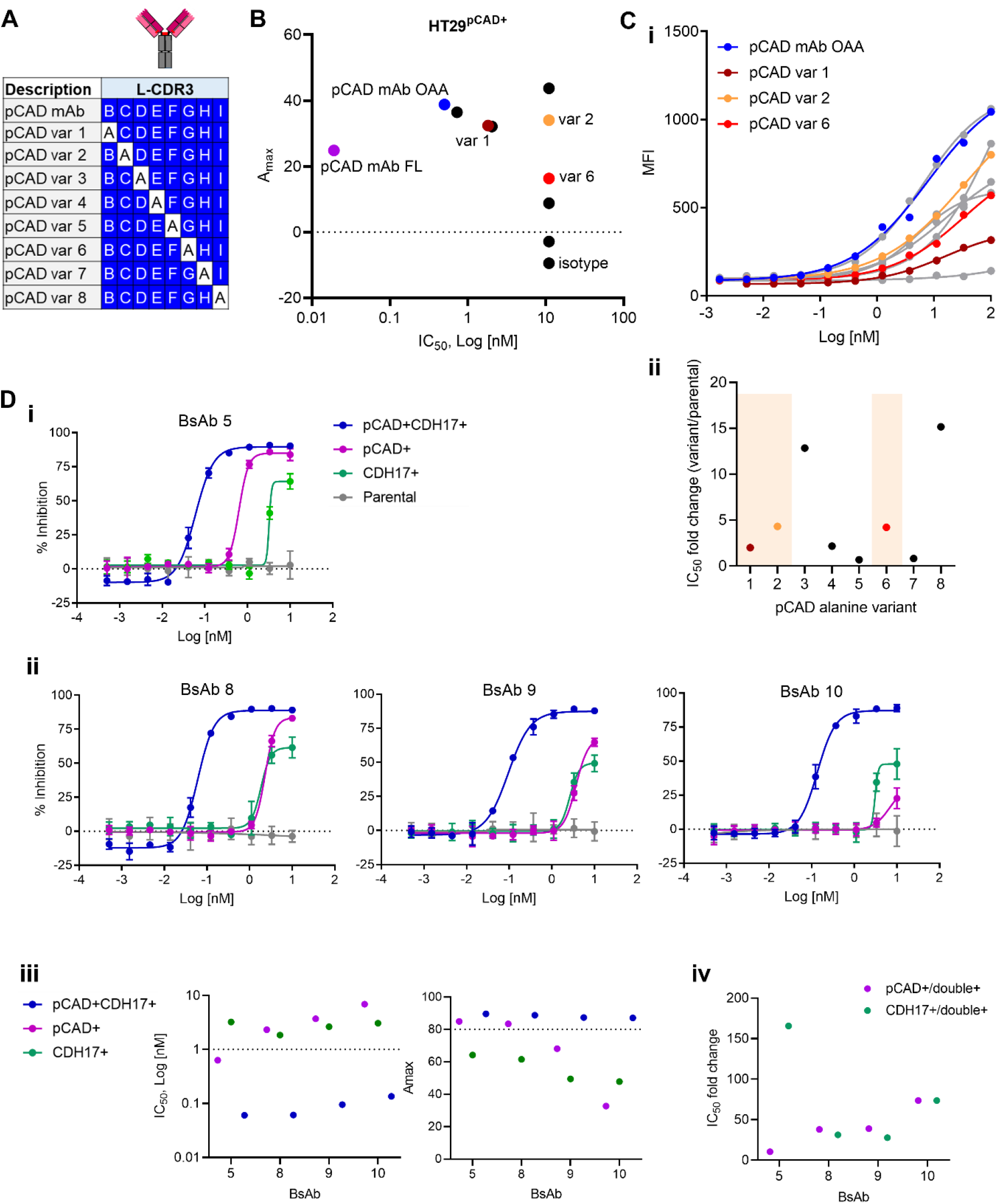
Selection of pCAD arm and generation of 2^nd^ round of bispecifics. (A) Representation of systematic alanine scanning of the light chain CDR3 to generate 8 alanine variants. (B) Internalization and proliferation inhibition assay performed using the HT29^pCAD+^ cell line treated with OAA pCAD alanine variants, and the control pCAD mAb (OAA and FL). Highlighted dots represent the 3 pCAD alanine variants that were selected for further investigation. (C) (i) FACS binding titration assay using the HT29^pCAD+^ cell line treated with OAA pCAD alanine variants, and the control pCAD mAb OAA. The 3 pCAD alanine variants selected are highlighted. (ii) Fold change of the IC_50_s between the pCAD alanine variants and parental pCAD. (D) Internalization and proliferation inhibition assay performed on a panel of HT29 cell lines treated with (i) the parental pCAD mAb x lead CDH17 bsAb (bsAb5), and (ii) the 3 pCAD alanine variants shuffled with the lead CDH17 arm (bsAb 8, 9, and 10). (iii) IC50 and Amax values and (iv) the IC_50_ fold change between inhibition of pCAD only and dual expressing cells, and CDH17 only and dual expressing cells, upon treatment with pCAD x CDH17 bsAbs.

Using the pCAD alanine variant OAAs, we ran an *in vitro* internalization and inhibition of cell proliferation assay in an engineered pCAD+ cell line (HT29^pCAD+^) **(Figure 3B)**. The calculated IC_50_ values ranged between 0.02 nM to 11 nM and the Amax values ranged between -9.5% to 62.7% inhibition, where all alanine variants achieved a less potent IC50 compared to the original pCAD mAb OAA. We also ran the pCAD alanine variant OAAs, along with pCAD mAb OAA, in a FACS binding assay using the same pCAD+ cell line **(Figure 3Ci)**. The IC_50_ values ranged between 0.23 nM to 100 nM, with most candidates achieving the desired decreased binding compared to pCAD mAb. Because the FACS binding assay provided a larger dynamic range for comparing pCAD alanine variants than the internalization and proliferation inhibition assay, we opted to prioritize the FACS binding results for pCAD arm selection. Taking the fold change of the IC50 values between the alanine variant and the parental pCAD arm, we selected those that had an intermediary of 1.5 to 5-fold less activity **(Figure 3Cii)**. From this list, we selected three variants that had low, intermediate, and high binding, pCAD var 1, var 6, and var 2, respectively, to combine with our lead CDH17 avidity-driven arm CDH17 11, to correspondingly generate bsAb 8, bsAb 9, and bsAb 10.

To evaluate the specificity of our three affinity detuned pCAD x CDH17 bsAbs, we ran all candidates in an *in vitro* internalization and inhibition of proliferation assay, across a panel of cell lines (HT29 parental with low expression of both antigens, HT29^pCAD+^, HT29^CDH17+^, and HT29^pCAD+CDH17+^) **(Figure 3D)**. As expected, across all bsAbs, no activity was observed in the parental double negative cell line, reduced activity was observed in the CDH17+ line, and high activity was observed in the dual-expressing cell line. Moreover, compared to the bsAb with the parental pCAD arm **(Figure 3Di)**, bsAb 10 was able to induce the largest reduction in activity in the pCAD-only cell line compared to bsAb 8 and bsAb 9 **(Figure 3Dii)**, as quantified by the increased IC50 and decreased Amax values **(Figure 3Diii)**, and fold change of IC_50_ values between single expressing cell lines and the dual expressing cell line **(Figure 3Div)**. Overall, in phase II, we generated a pCAD arm using an alanine scanning method that achieved specificity to dual expressing cells.

### pCAD x CDH17 bispecific ADC is selective for dual-expressing tumors in dual flank *in vivo* mouse models

To evaluate the *in vivo* antitumor activity and specificity of a pCAD x CDH17 bispecific ADC, we conjugated our lead pCAD x CDH17 bsAb (bsAb 10) with the cytotoxic payload monomethyl auristatin E (MMAE). MMAE is a potent antineoplastic agent that binds to tubulin and microtubules to induce cell death and is one of the most frequently employed ADC payloads in the clinic^26^. We also conjugated MMAE to the parental pCAD x CDH17 bsAb 5 and affinity detuned pCAD x CDH17 bsAb 8, which represents the selected alanine variant pCAD arm with lower cell binding affinity, to be used for comparative purposes. All three pCAD x CDH17 MMAE ADCs were tested *in vitro* using the HT29 panel previously used to confirm expected profiles **(Figure S4A)**.

For our *in vivo* experiments, we utilized two dual flank cell line-derived xenograft models to comprehensively evaluate the efficacy and specificity of the pCAD x CDH17 bispecific ADC against dual-antigen expressing tumors compared to single-antigen expressing tumors. One model was implanted with HT29 cells expressing both pCAD and CDH17 on the right flank, and with single positive HT29^CDH17+^ cells on the opposite flank, referred to as the CDH17 dual flank model **(Figure 4Ai)**. The second model was similarly implanted with HT29^pCAD+CDH17+^ cells on the right flank and single positive HT29^pCAD+^ cells on the opposite flank, referred to as the pCAD dual flank model **(Figure 4Aii)**. Both models’ expression of CDH17 and pCAD were evaluated and characterized using immunohistochemistry (IHC) in implanted engineered HT29 cell lines. At both 15- and 32-days post-implantation, CDH17 and pCAD were robustly expressed in the respective engineered xenografted tumors, as depicted in the representative images in **Figure S4B** at day 32. This confirms that the upregulation of CDH17 and pCAD in HT29 engineered cells is maintained throughout *in vivo* growth.

**Figure 4.**
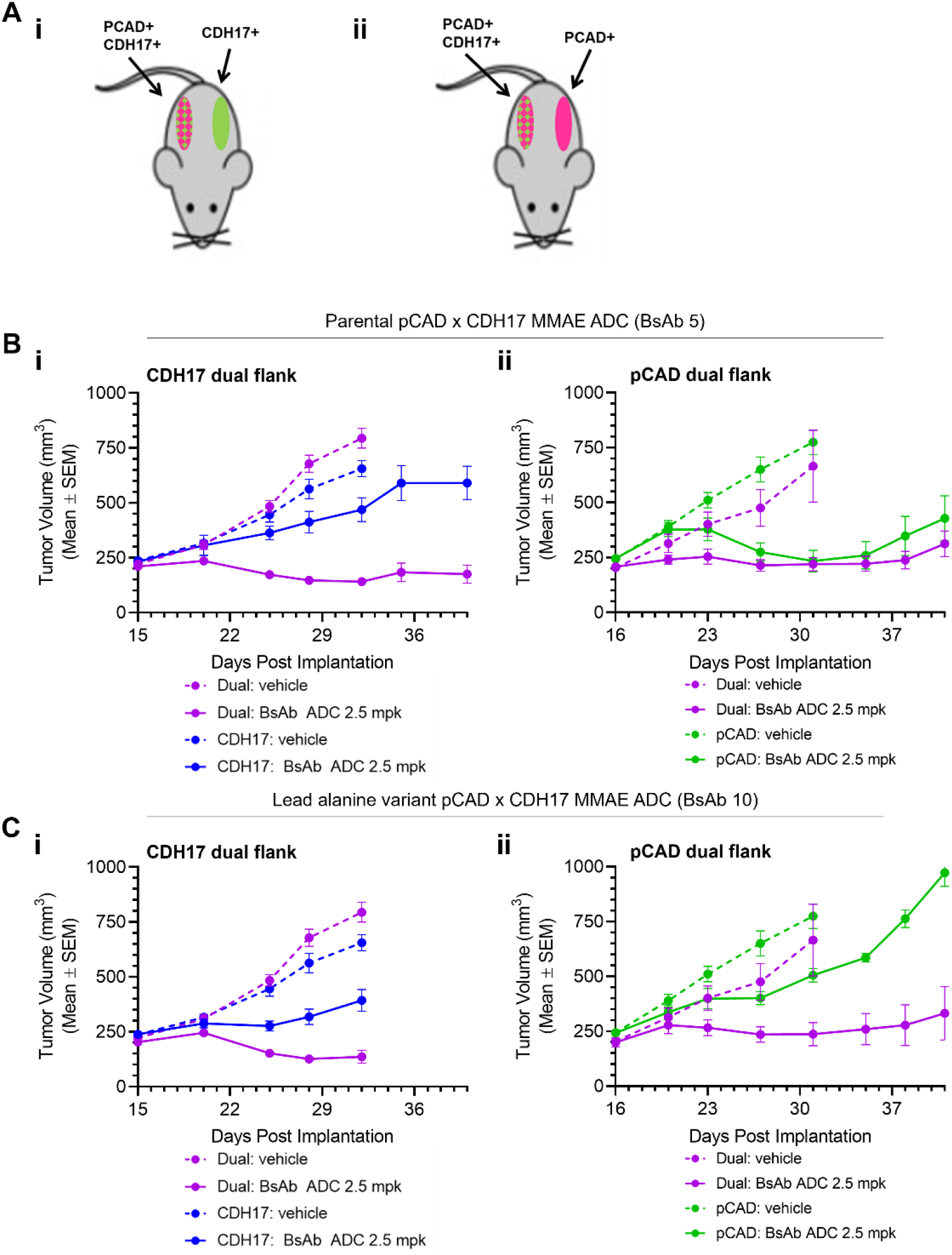
pCAD x CDH17 MMAE ADC has anti-tumor activity in dual flank mouse models. (A) Schematic of the two dual flank cell line-derived xenograft models used in these studies: (i) CDH17 dual flank model and (ii) pCAD dual flank model. (B) Treatment response in (i) CDH17 and (ii) pCAD dual flank models treated with the parental pCAD x CDH17 (bsAb 5) MMAE ADC at 2.5 mpk. Dotted lines represent vehicle treatment and solid lines represent treatment with the bsAb ADC. (C) Treatment response in both models as performed in (B), with the lead pCAD x CDH17 (bsAb 10) MMAE ADC at 2.5 mpk.

Upon administering a single intravenous dose of 2.5 mg/kg of the parental pCAD x CDH17 (bsAb 5) MMAE ADC, we observed expected *in vivo* responses that aligned with our *in vitro* observations. In the CDH17 dual flank model, the dual-antigen expressing tumor exhibited durable remission *in vivo*. Conversely, the CDH17-expressing tumor on the opposite flank continued to increase in tumor volume, demonstrating growth similar to that observed in the vehicle-treated CDH17-expressing tumor **(Figure 4Bi)**. Expectedly in the pCAD dual flank model, the single dose of the parental pCAD x CDH17 MMAE ADC lead to remissions in both the dual-antigen expressing tumor and the pCAD-expressing tumor, indicating non-specific targeting of the ADC **(Figure 4Bii)**.

Next, we treated both dual flank models with our lead alanine variant pCAD x CDH17 (bsAb 10) MMAE ADC. In the CDH17 dual flank model, we observed a trend consistent with the results of the parental pCAD x CDH17 MMAE ADC, as the CDH17 bsAb arm remained identical **(Figure 4Ci)**. However, the pCAD dual flank model exhibited a divergent response between the dual-antigen expressing tumor and the pCAD-expressing tumor upon treatment with bsAb 10 MMAE ADC, similar to our *in vitro* assay results **(Figure 4Cii)**. We further performed PK studies for measuring total antibody and ADC levels in the plasma and demonstrated similar exposure between the parental and affinity detuned ADCs up to 72 hours **(Figure S5A)**, with comparable total antibody and ADC half-lives (62 and 38 hours for bsAb 5 ADC; 77 and 40 hours for bsAb 10 ADC). Comparing the *in vivo* results between bsAb 5 MMAE ADC and the lead alanine variant bsAb 10 MMAE ADC confirms the successful affinity detuning of the pCAD arm, reducing tumor inhibition in pCAD+ single-antigen expressing tumors. Interestingly, when we treated both dual flank models with the alternative alanine variant pCAD x CDH17 (bsAb 8) MMAE ADC, we observed a difference in tumor responses exclusively in the CDH17 dual flank model **(Figure S5Bi)**. The pCAD+ tumor in the treated pCAD dual flank model reached a tumor volume similar to that of the dual-expressing tumor indicating non-specific dual targeting **(Figure S5Bii)**, as expected based on the *in vitro* internalization and proliferation inhibition assay results of bsAb 8 on HT29^pCAD+^ cells, underscoring the importance of the pCAD arm in driving selectivity. Our *in vivo* findings validate the increased specificity of our pCAD x CDH17 bispecific MMAE ADC towards dual-antigen expressing tumors, leading to selective and significant *in vivo* antitumor activity and achieving complete responses.

## Discussion

In the past several years, there has been a marked increase in ADCs entering clinical trials for solid tumors. ADCs hold immense potential in cancer therapy, but their clinical application faces challenges because of the need for highly potent payloads to effectively target and kill tumor cells. This necessity for payload potency underscores the importance of selecting target antigens that exhibit high and robust expression on tumor cells while being limited or absent on normal tissues, establishing a large therapeutic window. As a result, FDA-approved ADCs for solid cancers have predominantly focused on antigens like HER2, characterized by a substantial differential expression between tumor and normal cells^27^. Despite the emergence of a handful of clinically approved ADCs, the majority are HER2-targeting, demonstrating the challenge with identifying suitable antigens. To diversify our approach, we focused on dual targeting CDH17 and pCAD, both of which have been separately used as targeting antigens due to their prevalence in solid cancer tissues, such as CRC. Here, our hypothesis centered on addressing tumor specificity by requiring the co-expression of two surface cell antigens on tumor cells, bypassing potential normal tissue liabilities associated with each antigen when targeted individually, and for more specific delivery of the cytotoxic payload (MMAE). Other potential strategies to enhance tumor specificity include the selection of dual payloads that can elicit a synergistic effect, and more advanced payloads that target cancer cell-specific vulnerabilities.

In this study, we rationally selected for an avidity-driven CDH17 arm and pCAD arm by using antibodies in both a OAA and FL format, and selected those that necessitated two arms for high cell binding and payload delivery. This avidity-driven criteria resulted in a substantial increase in binding affinity, internalization and inhibition of proliferation efficiency, specifically in cells expressing both antigens compared to cells expressing either pCAD or CDH17 alone. Further, we demonstrated the ability to affinity detune pCAD arms, generating a broader diversity of antibodies. This expansion allowed us to identify arms with weaker binding affinities to pCAD+ expressing cells, while preserving their ability to internalize. These arms were subsequently coupled to CDH17 and tested to determine the optimal pairing that achieved a significant difference in response between dual-expressing and pCAD-only expressing cell lines. Ultimately an intermediate cell binding candidate emerged as the ideal option, that maintained sufficient, but less binding compared to the parental pCAD arm. The suitability of the lead BsAb was confirmed when conjugated to MMAE and tested *in vivo* with two dual flank models, where it exhibited a high degree of pCAD and CDH17 dual-positive tumor specificity, highlighting the effectiveness of our rational bsAb design and selection process. This unique avidity-driven approach can be implemented to other co-expressing tumor antigens that are being considered for dual-specific targeting.

While this study confirms the widespread presence of both pCAD and CDH17 cell surface expression on CRC cells, challenges with tumor heterogeneity remain. Certain tumors exhibit a higher proportion of cells that co-express both antigens, while others predominantly consist of either pCAD or CDH17 single-expressing tumor cells resulting in a small proportion of dual positive cells. A well-known phenomenon associated with ADCs is the bystander effect, where the released payload crosses the cell membrane following lysosomal degradation of the ADC and enters neighboring cells to exert its cytotoxic effects beyond the initially targeted cell. Therefore, evaluating the bystander effect with our pCAD x CDH17 MMAE ADC would not only provide insight into the extent by which neighboring cells can be affected, but also elucidate the ratio of dual-expressing cells required to induce significant damage across the whole tumor. Understanding these dynamics would aid in addressing the complexities of tumor heterogeneity that minimizes a large-scale effect across tumors, and thus be important for optimizing the therapeutic potential of a pCAD x CDH17 MMAE ADC in CRC.

In summary, our preclinical investigation demonstrates a promising avenue for utilizing pCAD and CDH17 as dual targeting antigens in the context of ADCs for CRC. Through a systematic screening process involving CDH17 arms and affinity-detuned pCAD arms, we identified and selected monovalent arms that exhibited optimal properties when paired as a bispecific ADC. The preclinical data here of a pCAD x CDH17 bispecific MMAE ADC warrants further investigation in the pursuit of improved treatment options for CRC patients.

## Materials & Methods

### Tabula Sapiens scRNAseq Dataset Processing

Raw gene expression count of 454,069 cells from 15 normal human subsets across 24 organs were downloaded from the Tabula Sapiens Consortium^23^. The original cell type labeling was inherited. Pseudobulk counts were created by adding gene expression counts in cells of the same cell type and from the same patient. Trimmed mean of M values (TMM) normalization from the edgeR package followed by counts per million (CPM) normalization were applied to remove donor and cell type size effect.

### Public CRC scRNAseq Dataset Processing

Raw count from two public CRC scRNAseq datasets were downloaded from Jabbari et al.^24^ and Qian et al.^25^. The SCTransform function from the Seurat package^28^ was applied for normalization before the Symphony package^29^ was leveraged for automatic cell type annotation referencing our internal immune-oncology scRNAseq-profiled cell types. Pseudobulk counts were created by adding gene expression counts in cells of the same predicted cell type and from the same patient. TMM normalization from the edgeR package followed by counts per million (CPM) normalization were applied.

### Cell Culture

Snu16 (ATCC) was cultured in RPMI 1640 (Gibco) supplemented with 10% FBS (Gibco). HT29 CDH3 knockout cell line was made with a viral vector that expressed Cas9 and a pCAD guide RNA. For engineering overexpressing cell lines, HT29 cell lines were transduced with a lentivirus construct expressing hpCAD, hCDH17, or hpCAD and hCDH17 under an EF1A promoter and were cultured in

McCoy’s 5A media (Gibco) supplemented with 10% FBS (Gibco). All cells were cultured at 37°C under 5% CO_2_ and passaged before reaching confluency up to 25 passages. Cells were validated for pCAD and/or CDH17 expression in FACS or were tested for sensitivity to CDH17+pCAD+ ADCs in a cytotoxicity assay prior to use.

### Internalization and Inhibition of Cell Proliferation Assay

Cells (Snu16, HT29 pCAD KO, HT29 pCAD+, HT29 CDH17+ or HT29 pCAD+CDH17+) were seeded in a culture-treated 384-well clear plate (2000 cells per well in 40 μL culture medium) and incubated at 37°C under 5% CO_2_ for 24 hours. Serially diluted samples (5 μL) were added to each well and a final concentration of 34 nM of anti-human Fc-VC-MMAE ADC (DAR 2.0) (5 uL) was added to each well. The plate was incubated at 37°C for 120 hours, then 25 uL of CellTiter-Glo (Promega) was added to each well, incubated at 25°C for 10 minutes, followed by measuring luminescence using the PHERAstar FSX Microplate reader. IC50 values were calculated using Graph Pad Prism 8 software. All assays were performed in quadruplicate technical replicates.

### Flow Cytometry Binding Assay

Binding to tool cell lines (HT29 pCAD KO, HT29 pCAD+, HT29 CDH17+, and HT29 pCAD+CDH17+ cells) was evaluated using fluorescence-activated cell sorting (FACS). All antibodies were diluted in FACS buffer, and added to cells followed by an 1 hour incubation. Unbound antibodies were washed off, and cells were then incubated with Alexa Fluor® 647 AffiniPure Goat Anti-Human IgG (H+L) (1:250) (Jackson). After a 30 min incubation, unbound secondary antibody was washed off. Cells were then resuspended in FACS buffer with DAPI (1:5000) (Invitrogen) for FACS detection using the Milltenyi MACSQuant. The mean fluorescence intensity of the cells in the live gate was plotted against log antibody concentration, and the IC50 was determined by nonlinear regression fitting.

### Antibodies

All antibodies were cloned into CMV-promoter-driven expression plasmids for mammalian cell expression and subsequently produced by transient transfection in the HEK-293T cells. For the identification of avidity-driven antibodies, full monoclonal antibodies were produced as wild type huIgG1. The monovalent one-armed antibodies were produced using the knobs-into-holes technology (Merchant, 1998). The full length antibody heavy chain was modified to encode the T366W: S354C mutations, and the Fc fragment was modified to contain the Y349C:T366S:L368A:Y407V mutations. The light chain, mutated heavy chain, and Fc fragment were co-transfected into HEK-293 cells. Purification was done by affinity chromatography on a mAb SelectSure resin (Cytiva) followed by a polishing step by size exclusion chromatography. Antibodies were characterized for integrity and homogeneity by SDS-PAGE analysis and mass spectrometry.

The bispecific antibodies were also produced using the knobs-into-holes technology. The anti-pCAD antibodies harbor a Kappa light chain and the anti-CDH17 antibodies harbor a lambda light chain. The anti-pCAD heavy chain was modified to encode the T366W: S354C mutations, and the anti-CDH17 heavy chain was modified to contain the Y349C:T366S:L368A:Y407V mutations. The conjugated bispecific antibodies evaluated *in vivo* were designed with mutations E152C:S375C for facilitating conjugation and mutations D265A:P329A for silencing Fc effector functions. The purification of bispecific antibodies was done by affinity chromatography on a mAb SelectSure resin (Cytiva) followed by a first polishing step by affinity chromatography on a CaptureSelect LC Lambda resin (Thermofisher) and a second polishing step by size exclusion chromatography. Antibodies were characterized for integrity and homogeneity by SDS-PAGE analysis and mass spectrometry.

Other antibodies used in this study were purchased from commercial vendors: Alexa Fluor® 647 AffiniPure Goat Anti-Human IgG (H+L) (Jackson), anti-CDH17 (abcam 109190), anti-pCAD (Sigma HPA001767).

### BsAb Conjugation

Ten mg of each antibody was incubated with RMP Protein A resin (GE) at a ratio of 10 mg Ab to 1 mL resin in PBS for 15 minutes with mixing in an appropriately sized disposable column. Cysteine HCl was added to a final concentration of 20 mM and incubated with agitation for 30 mins at room temperature to allow the reactive cysteines to be deblocked. The resin was quickly washed with 50 column volumes PBS on a vacuum manifold. The resin was then re-suspended in an equal volume PBS containing 250 nM CuCl2. Reformation of antibody interchain disulfides was monitored by taking time points. At each time point, 25 mL of resin slurry was removed, 1 mL of 20 mM MC-VC-MMAE was added, and the tube flicked several times. The resin was spun down, supernatant removed, and then eluted with 50 mL antibody elution buffer (Thermo). The resin was pelleted and the supernatant analyzed by reverse phase chromatography using an Agilent PLRP-S 4000A.

Once it was determined that the antibody reformed its interchain disulfide bonds, the resin was washed with 10 column volumes PBS and the resin was resuspended in an equal volume PBS and 8 equivalents of MC-VC-MMAE in DMSO, with a final concentration of 10% DMSO in the reaction and then incubated at room temperature for 3 hours. The resin was then washed with 50 column volumes PBS. The ADC was eluted from the protein A resin with antibody elution buffer. The ADC was then buffer exchanged into PBS. Samples were sterile filtered with a 0.22 mm syringe filter. The following analyses were performed -analytical SEC to determine percent monomer, mass spectroscopy to determine DAR, LAL test to determine endotoxin load and protein concentration was determined by A280 utilizing extinction coefficient and molecular weight of antibody or BCA protein assay (Thermo).

### In Vivo *Dual Flank Model Efficacy Studies*

Studies were conducted in accordance with ethical guidelines and regulations set by the Novartis BioMedical Research Institutional Animal Care and Use Committee, as well as the Guide for the Care and Use of Laboratory Animals. Cell lines were confirmed to be free of mycoplasma and mouse viruses before use and cultured in medium according to ATCC guidelines. 5x10E6 cells were resuspended in 200 uL of ice-cold Hank’s Balanced Salt Solution (HBSS; Gibco, catalog # 25200-056) containing 50% Matrigel (Corning, catalog #354234). Female NSG mice (6-12 weeks of age) were inoculated subcutaneously in the right flank with HT29^pCAD+CDH17^+ cell line and concurrently inoculated on the left flank with either the HT29^pCAD+^ or the HT29^CDH17+^ cell line. Tumor bearing mice were randomized into treatment groups 15 or 16 days after inoculation, once average tumor volumes on both flanks reached 200-225 mm^3^. Tumors were measured twice weekly by calipering in two dimensions. Tumor volume was calculated using a modified ellipsoid formula: tumor volume -L × W2 × π/6, where L is the longest axis of the tumor and W is perpendicular to L. Serum samples were collected 1 hour, 24 hours, and 72 hours post-dose for quantification of total monoclonal antibodies (mAb) and total ADC-MMAE concentrations.

## Notes

### Competing Interest Statement

All authors of this manuscript have been or currently are employees of Novartis Pharma AG during the time of preparation.

