## Supplemental Figures for "A bispecific antibody-drug conjugate targeting pCAD and CDH17 has antitumor activity and improved tumor specificity"

[1]Novartis Biomedical Research, Cambridge, MA, USA.

[2]Novartis Biomedical Research, Basel, Switzerland.

[#]These authors contributed equally.

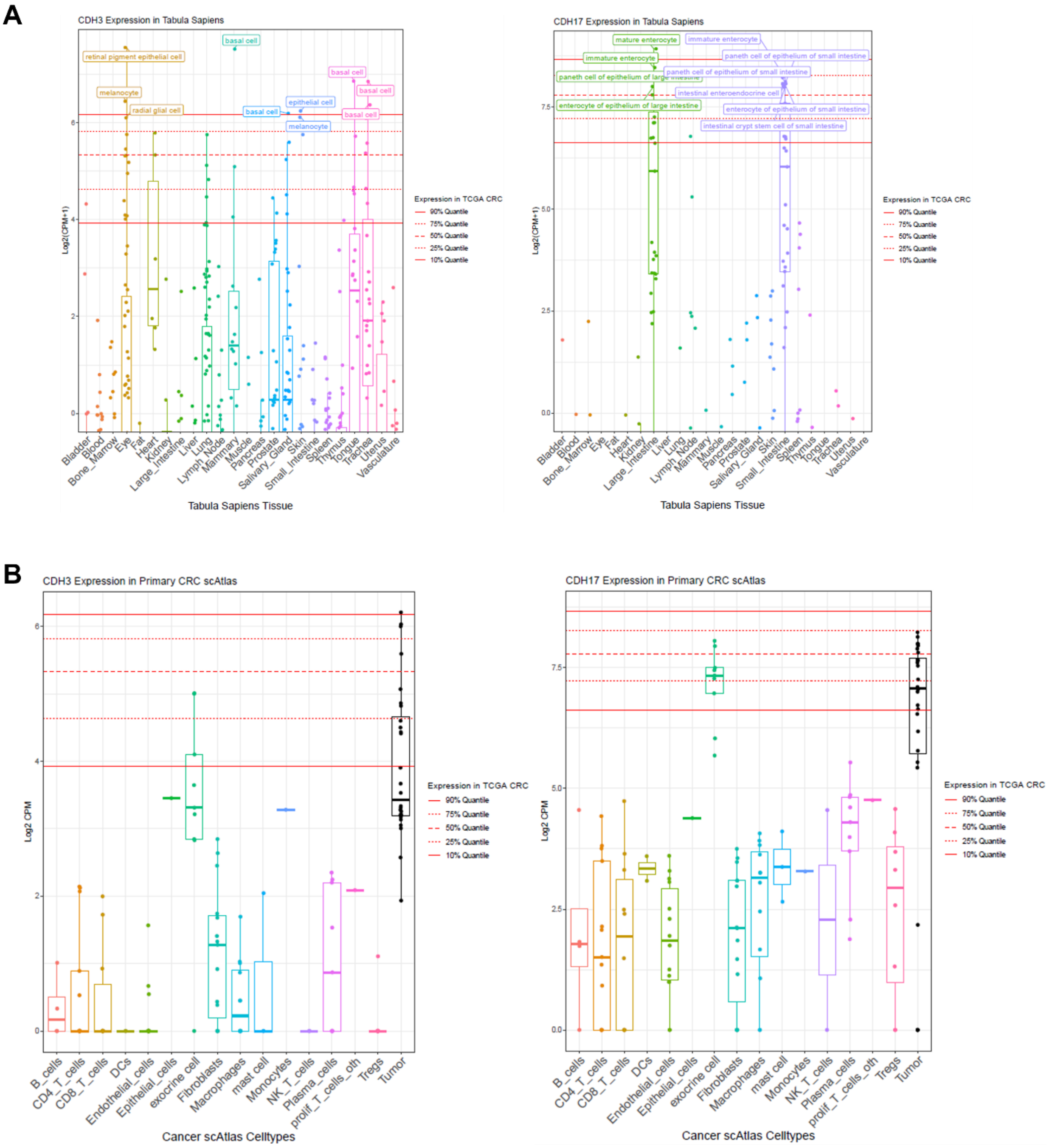


**SI Figure 1. Single cell RNA sequencing datasets for *CDH3* and *CDH17* expression.** (A) scRNAseq data from healthy normal donors demonstrating distributed *CDH3* single cell RNA expression, and isolated *CDH17* single cell RNA expression in the small and large intestine. (B) scRNAseq data from CRC patients confirm that cancer cells, rather than other cell types in the tumor microenvironment, are responsible for high *CDH17* and *CDH3* expression.

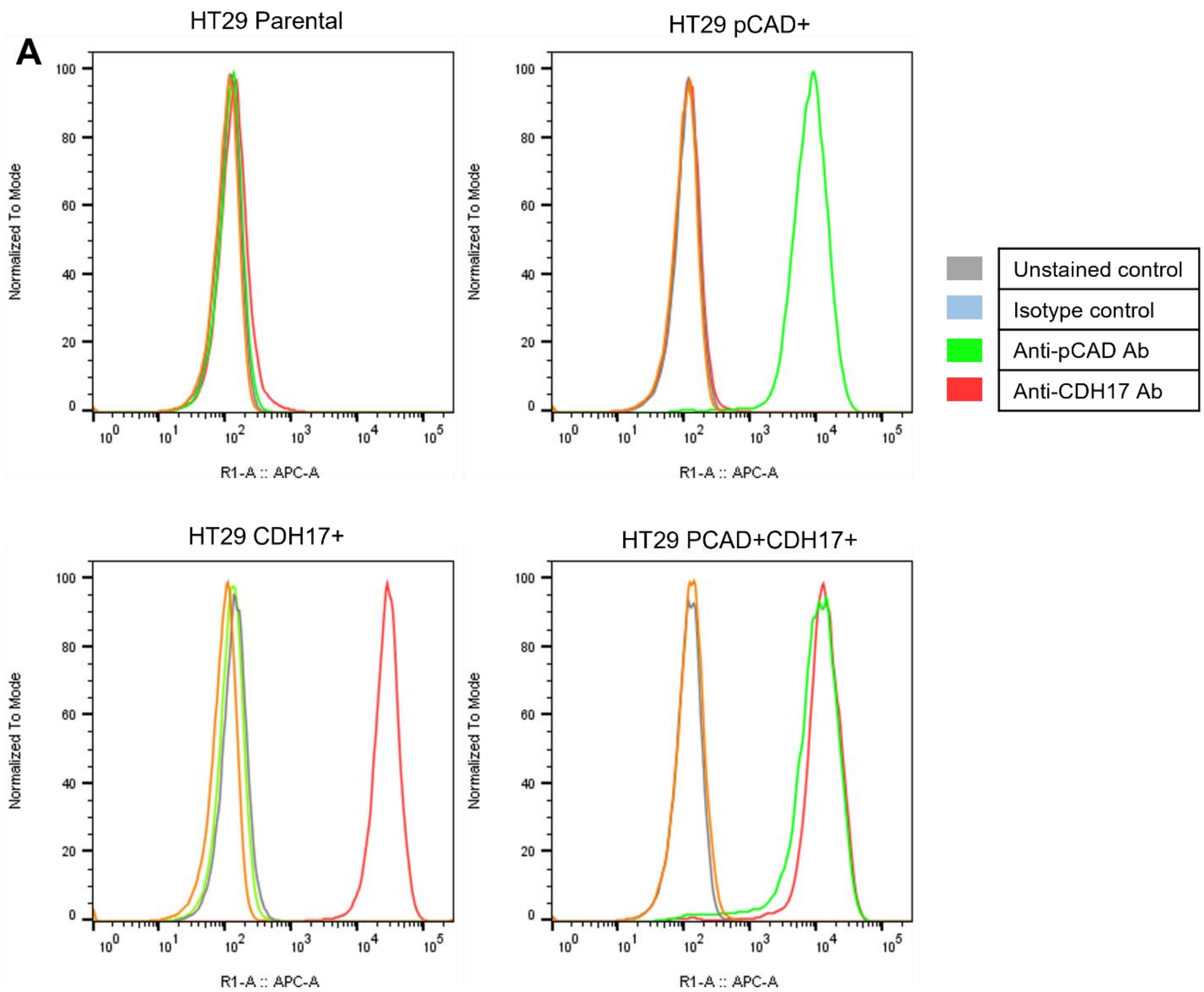


**SI Figure 2. Using engineered CRC cell line HT29 as a tool model.** (A) FACS profiling of the HT29 parental line with endogenous low expression of pCAD and CDH17, and after engineering pCAD-overexpressing ($HT29^{pCAD+}$), CDH17-overexpressing ($HT29^{CDH17+}$) , and double positive pCAD- and CDH17-overexpressing ($HT29^{pCAD+CDH17+}$) cell lines in the HT29 background.

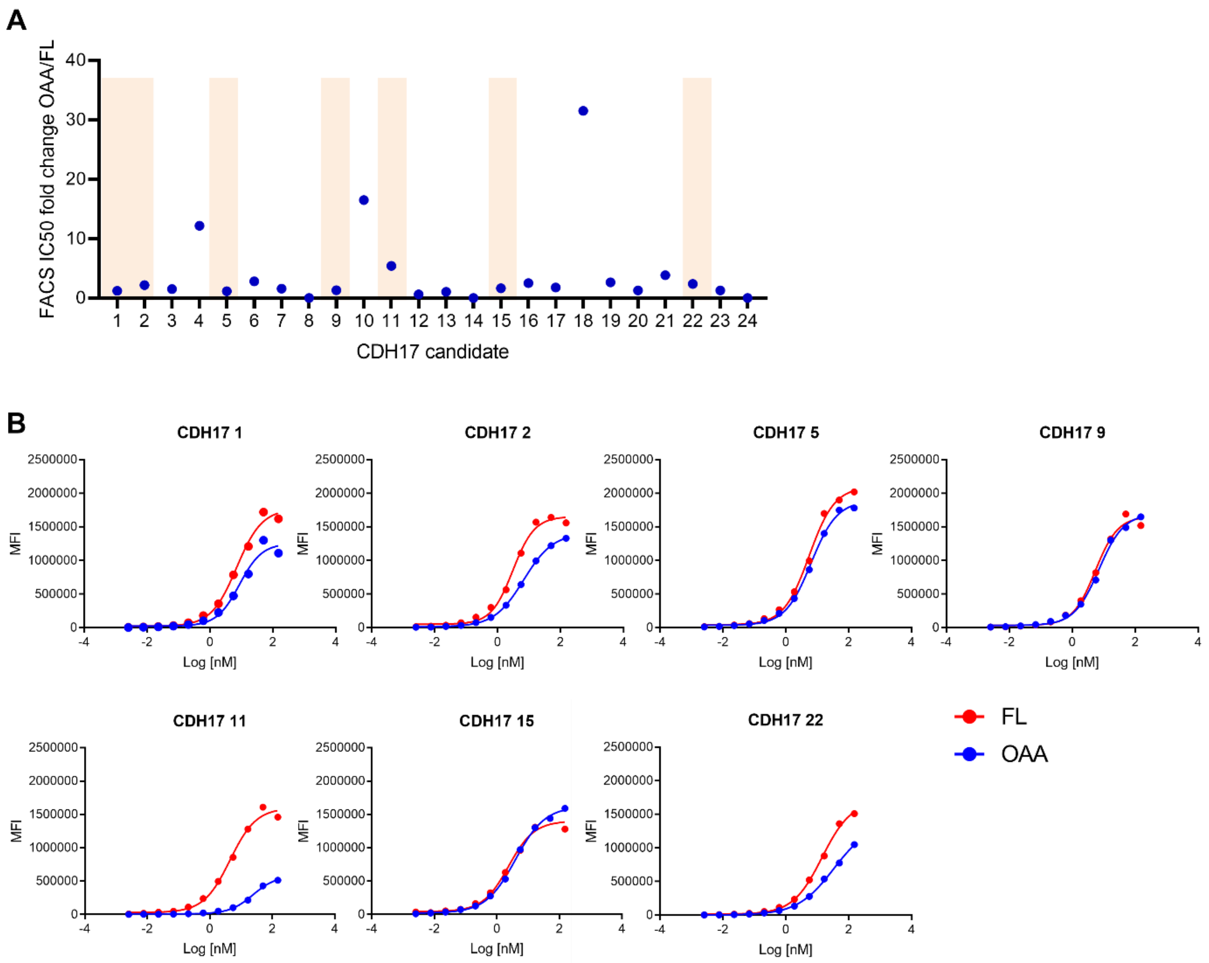


**SI Figure 3. FACS binding of CDH17 OAA and FL antibodies.** (A) Fold change between the IC50 of OAA binding and the IC50 of FL binding. (B) FACS binding titration curves of the 7 CDH17 arms selected based on the results from the *in vitro* internalization and inhibition of proliferation assay. Across all 7 CDH17 candidates, the OAAs had the same or less binding compared to the FL antibody.

**A** **i**

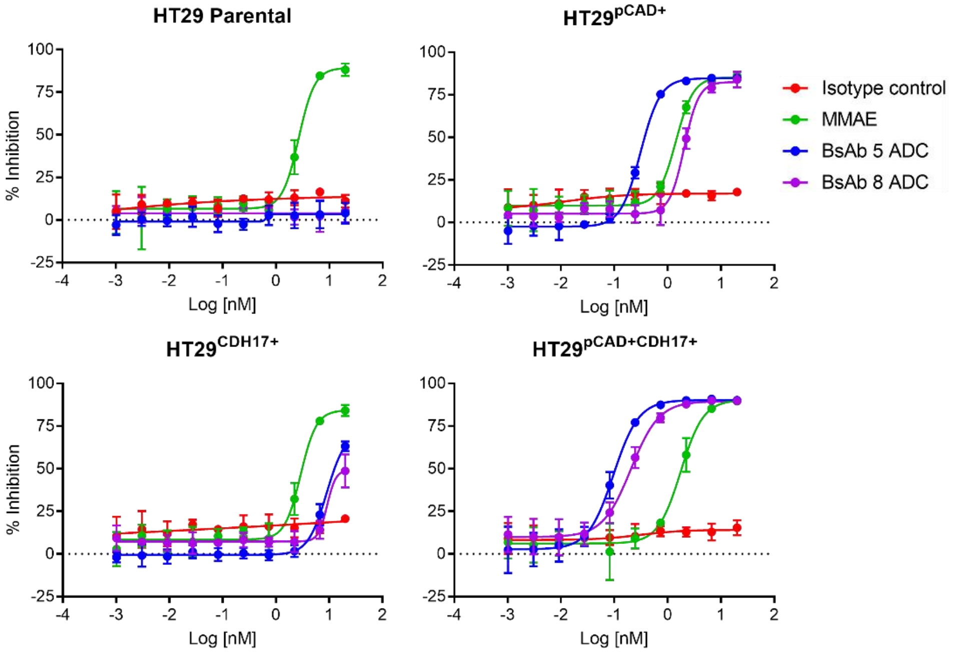


**ii**

| **IC50 (nM)** | Isotype control | MMAE | BsAb 5 ADC | BsAb 8 ADC |
|---|---|---|---|---|
| HT29 Parental | - | 2.67 | - | - |
| HT29 pCAD+ | - | 1.43 | 0.31 | 2.05 |
| HT29 CDH17+ | - | - | - | - |
| HT29 pCAD+CDH17+ | - | 1.76 | 0.10 | 0.21 |

**B** **i**

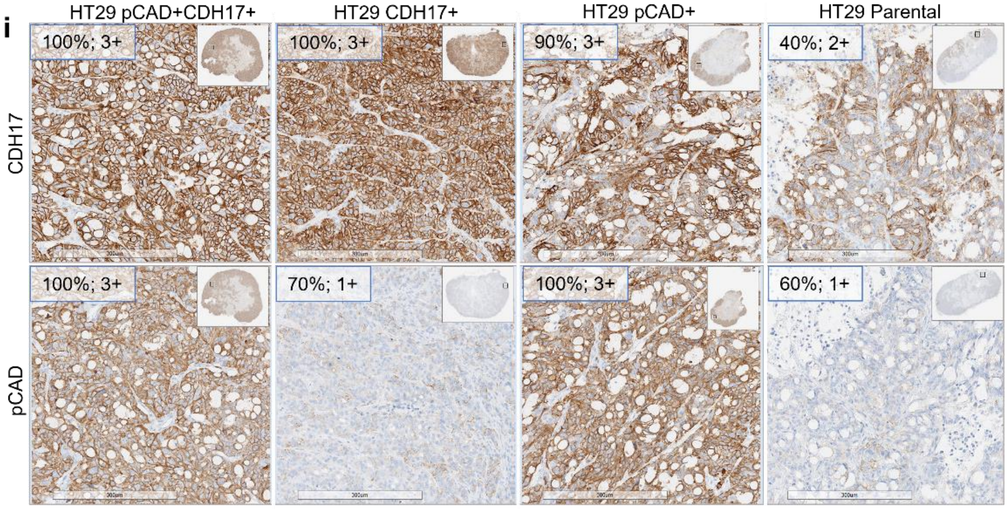


**SI Figure 4. Validation of *in vivo* material (pCAD x CDH17 MMAE ADCs) and *in vivo* IHC profiling for dual flank mouse model development.** (A) (i) *In vitro* validation of bsAb MMAE ADC material in panel of HT29 cells before *in vivo* experiment, and (ii) IC50 values. (B) IHC characterization at 32 days post-implantation of HT29 cells demonstrate that the upregulation of CDH17 and pCAD in HT29 engineered cells is maintained throughout *in vivo* growth. Notably, CDH17 levels remained detectable in both the opposite engineered and parental HT29 xenografts.

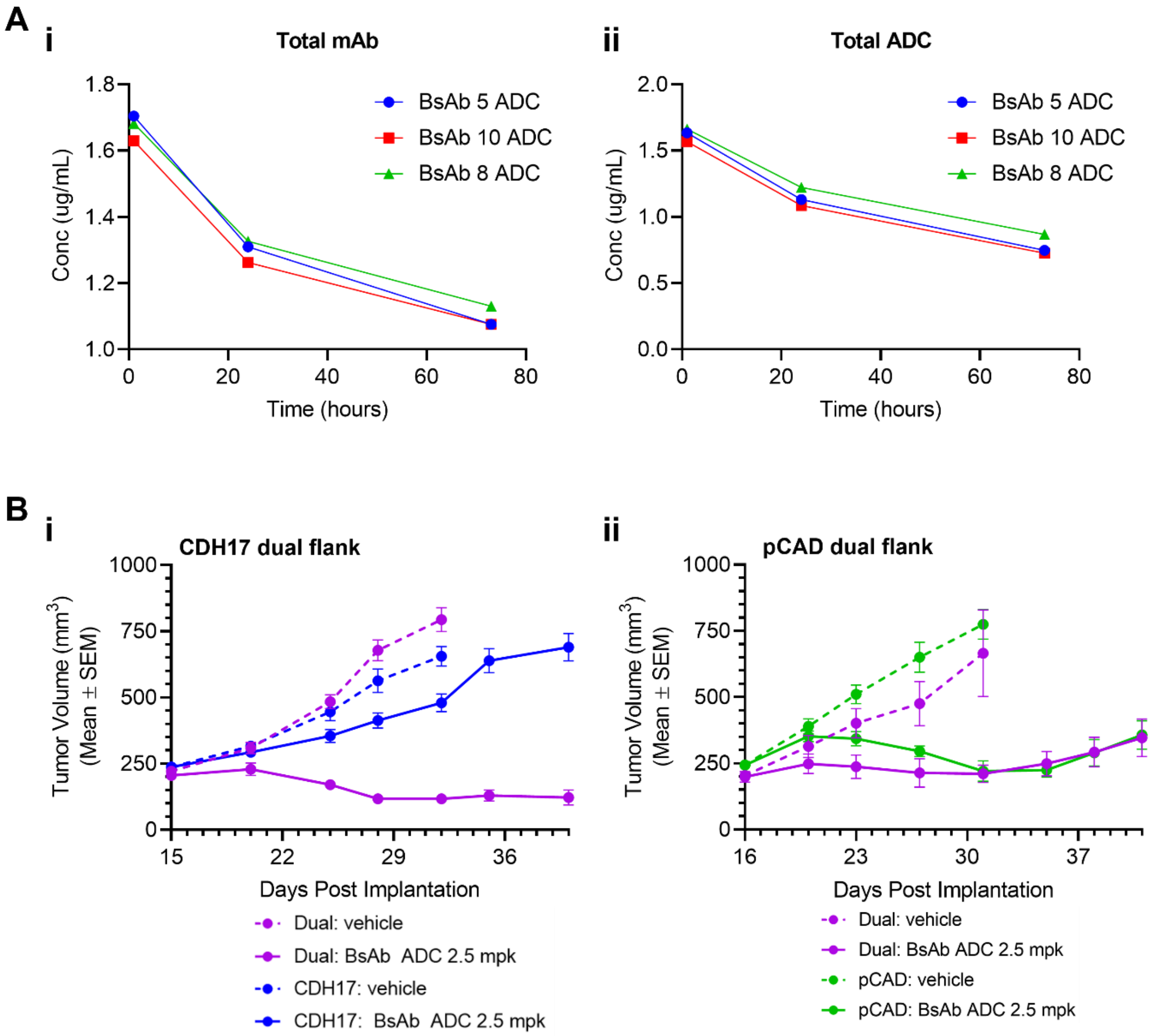


**SI Figure 5. Additional *in vivo* bsAb MMAE ADC results.** (A) PK study measuring (i) total antibody and (ii) total ADC levels in plasma, demonstrating similar exposure across all three ADCs. (B) Treatment response in (i) CDH17 and (ii) pCAD dual flank models treated with the alternative alanine variant pCAD x CDH17 (bsAb 8) MMAE ADC at 2.5 mpk.
